# Oviposition and vertical dispersal of *Aedes* species Meigen 1818 (Diptera: Culicidae) at different heights and seasonal periods in an urban forest fragment in Manaus, Amazonas, Brazil

**DOI:** 10.1101/2024.05.09.593399

**Authors:** William Ribeiro da Silva, Adriano Nobre Arcos, Francisco Augusto da Silva Ferreira, Joelma Soares-da-Silva, Grafe Oliveira Pontes, Mário Antônio Navarro da Silva, Rosemary Aparecida Roque, João Antonio Cyrino Zequi

**Affiliations:** Programa de Pós-Graduação em Ciências Biológicas (Entomologia), Instituto Nacional de Pesquisas da Amazônia (INPA), Avenida André Araújo, 2936, Petrópolis, Manaus, Amazonas, 69067-375, Brasil; Laboratório de Hidrologia, Instituto Nacional de Pesquisas da Amazônia (INPA), Avenida André Araújo, 2936, Petrópolis, Manaus, Amazonas, 69067-375, Brasil; Núcleo Patógenos, Reservatórios e Vetores na Amazônia (PreV Amazônia) – FIOCRUZ Amazônia; Curso de Ciências Naturais, Campus VII, Universidade Federal do Maranhão (UFMA), Avenida Dr. José Anselmo, 2008, São Sebastião, Codó, Maranhão, 65400-000, Brasil; Unidade de Entomologia Nelson Ferreira Fé - UENFF, Fundação de Medicina Tropical Doutor Heitor Vieira Dourado (FMT – HVD), Av. Pedro Teixeira, Dom Pedro, Manaus, Amazonas, 69040-000, Brasil; Laboratório de Morfologia e Fisiologia de Culicidae e Chironomidae, Universidade Federal do Paraná (UFPR), Caixa Postal 19020, Curitiba, PR, Brasil; Laboratório de Malária e Dengue, Instituto Nacional de Pesquisas da Amazônia, Avenida André Araújo, 2936, Petrópolis, Manaus, Amazonas, 69067-375, Brasil; Laboratório de Entomologia Médica, Departamento de Biologia Animal e Vegetal, Universidade Estadual de Londrina (UEL). Caixa Postal 6001, 86051-970 Londrina, Paraná, Brasil

**Keywords:** Vertical dispersal, dengue, Zika, *Aedes aegypti*

## Abstract

Mosquitoes of the genus *Aedes* stand out for their high susceptibility to several groups of arboviruses, especially those that cause dengue fever, Zika, and Chikungunya fever. However, aspects related to the vertical distribution of species in large urban centers are still poorly understood, therefore, this study aims to evaluate the dispersal and oviposition of *Aedes* at different height levels and seasonal periods. The study was developed in a tower with six floors located in an urban forest fragment, measuring 15.13 meters (m) high and 3.20 meters at the base. The following height ranges were considered: ground: 0 m; 1.20 m; 2.50 m; 3.60m; 4.90 m; 6m; 7.30m; 8.40m; 9.70 m; and 10.8 m. Three ovitraps were installed on each floor, separated by a distance of 1.50 m, totaling 30 for each sampling period. The ovitrap positivity index (OPI) and egg density index (EDI) were evaluated in order to monitor *Aedes* populations in different height ranges and also in different seasonal periods. The data demonstrated that lower heights show a greater abundance of *Aedes* eggs, however, this variable did not prove to be a limiting factor for mosquito colonization at the other heights evaluated. Furthermore, climatic factors, such as relative humidity have a positive influence (p<0.05) on the average number of eggs in the urban area of Manaus, especially during the dry period. These findings demonstrate that the vertical growth of urban centers can act positively tin increasing the density of *Aedes* and can influence the incidence of dengue and other arboviruses.

## INTRODUCTION

Culicidae (Diptera: Culicidae) are popularly known as mosquitoes, muriçocas, and carapanãs, coming from two subfamilies (Anophelinae and Culicinae) that include species of importance for global public health. Most of these insects occur in the five biogeographic regions of the globe with tropical forest environments, which hold the greatest diversity of species (Harbach, 2023; WRBU, 2023).

The genus *Aedes* Meigen 1818 includes species of medical and veterinary importance with a wide and growing global distribution (Iwamura et al. 2020). *Aedes aegypti* (Linnaeus, 1762) and *Aedes albopictus* (Skuse, 1895) are considered the most important species in the group, due to their implications for the transmission of many pathogens, such as the viruses that cause dengue, Zika, chikungunya, and urban yellow fever (Vega-Rúa *et al*. 2014; Campbell *et al*. 2015; Marcondes and Ximenes 2016; Leta *et al*. 2018).

Native to Africa, *A. aegypti* is found on other continents, mainly in tropical and subtropical regions (Powell *et al*. 2018), as well as in temperate regions, with its area predicted to expand by 2 to 6 km/year until 2050 (Iwamura et al. 2020). On the other hand, *A. albopictus* comes from Asia, and is present on the islands of the Western Pacific, the Indian Ocean, and on all continents, except Antarctica (Bonizzoni *et al*. 2013; Kraemer *et al*. 2015; Kamal *et al*. 2018).

Some environmental and anthropic factors were decisive for the propagation and permanence of species in these environments (Dickens *et al*. 2018). Furthermore, the ability of eggs to resist adverse conditions for a long period, due to the process called quiescence, is a fundamental aspect for the dispersal of these mosquitoes that carry different etiological agents (Soares-Pinheiro *et al*. 2017).

*A. aegypti* is predominant in the urban perimeter, where it is constantly associated with the presence of people, using different breeding sites that enable its propagation and maintenance (Carvalho and Moreira 2017). *A*. *albopictus* is beginning to enter this habitat, coexisting with the former species, however, its prevalence is in areas with native and/or secondary forest close to the urban environment (Forattini, 2002; Ferreira-de-Lima *et al*. 2020). This species could be a key vector in urban forest fragments, implying the main routes for the proliferation of pathogens between urban and wild environments, which have constant movement of people and animals (Montagner *et al*. 2018; Hendy *et al*. 2020).

Females of these species have similar oviposition strategies, distributing eggs in installments in breeding sites (Clements, 1992; Forattini, 2002) to ensure reproductive success (Carvalho and Moreira 2017; Ferreira-de-Lima et al. 2020). In general, breeding sites present a diversity of combinations (Soares-da-Silva et al. 2012; Goddard et al. 2017; González et al. 2020) and chemical parameters to help in the oviposition process (Abreu et al. 2015; Sanoussi et al. 2015; Hashim et al. 2018). Likewise, the presence of conspecific and heterospecific larvae are fundamental aspects in this process (Gonzalez et al. 2015).

Therefore, the choice of the ideal place for the eggs to adhere depends on multiple factors. In this context, ovitraps are potential breeding sites, as they provide extremely suitable conditions for oviposition and egg adherence (Depoli et al. 2016). These traps, when accompanied by grass infusion (attractant solution), become more inviting for females to lay eggs and, consequently, enable monitoring and control, and they end up removing eggs from the environment (Silva et al. 2018; Rossi-da-Silva et al. 2021).

Another important aspect that can also influence the oviposition process, specifically the abundance of eggs, are environmental variables (Custódio et al. 2019). The increase in precipitation frequency results in an increase in mosquito density due to the formation of new breeding sites, however, when extremely high, it can cause an abundance of these arthropods (Santos et al. 2020). On the other hand, high values of temperature and relative humidity can negatively affect different aspects of insect biology (Couret et al. 2014).

Mosquito dispersal is also affected by the availability of breeding sites maintained at ground level (Liew and Curtis 2004; Bergero et al. 2013; Ayllón et al. 2018) and vertically (Lau et al. 2013; Roslan et al. 2013; Jayathilake et al. 2015; Hamid et al. 2020). This “jumping oviposition” behavior is a strategy to avoid competition from larvae for food and ensure the reduction in risks associated with temporary breeding sites, making it possible to achieve reproductive success for the species (Clements, 1992; Forattini, 2002; Carvalho e Moreira 2017; Ferreira-de-Lima *et al*. 2020).

Given the importance of the oviposition process for the dispersal and maintenance of mosquitoes in different environments, there is still a need to carry out research related to the vertical dispersal of mosquitoes of the genus *Aedes*. Understanding the basic aspects of mosquito biology in the oviposition process, as well as verifying the effect of environmental variables on the abundance of eggs, is essential to carry out monitoring and control more efficiently. Therefore, the present study analyzed the frequency of *Aedes* oviposition at different heights and different seasonal periods, as well as verifying the effect of environmental variables (temperature, relative humidity, and precipitation) on the abundance of eggs.

## MATERIAL AND METHODS

### Study area

The study was carried out in a water tower restricted to an urban forest fragment, located on Campus I of the National Amazon Research Institute (INPA) (Figure 1). The institution is located in the municipality of Manaus/Amazonas, located in the center of the largest tropical forest in the world, with a humid equatorial climate, an average annual temperature of 26.7°C, and relative humidity of around 70%. The region has two well-defined seasons: rainy (December to June) and dry (July to November), based on precipitation and river levels (Lemos and Costa, 2017).

**Figure 1.**
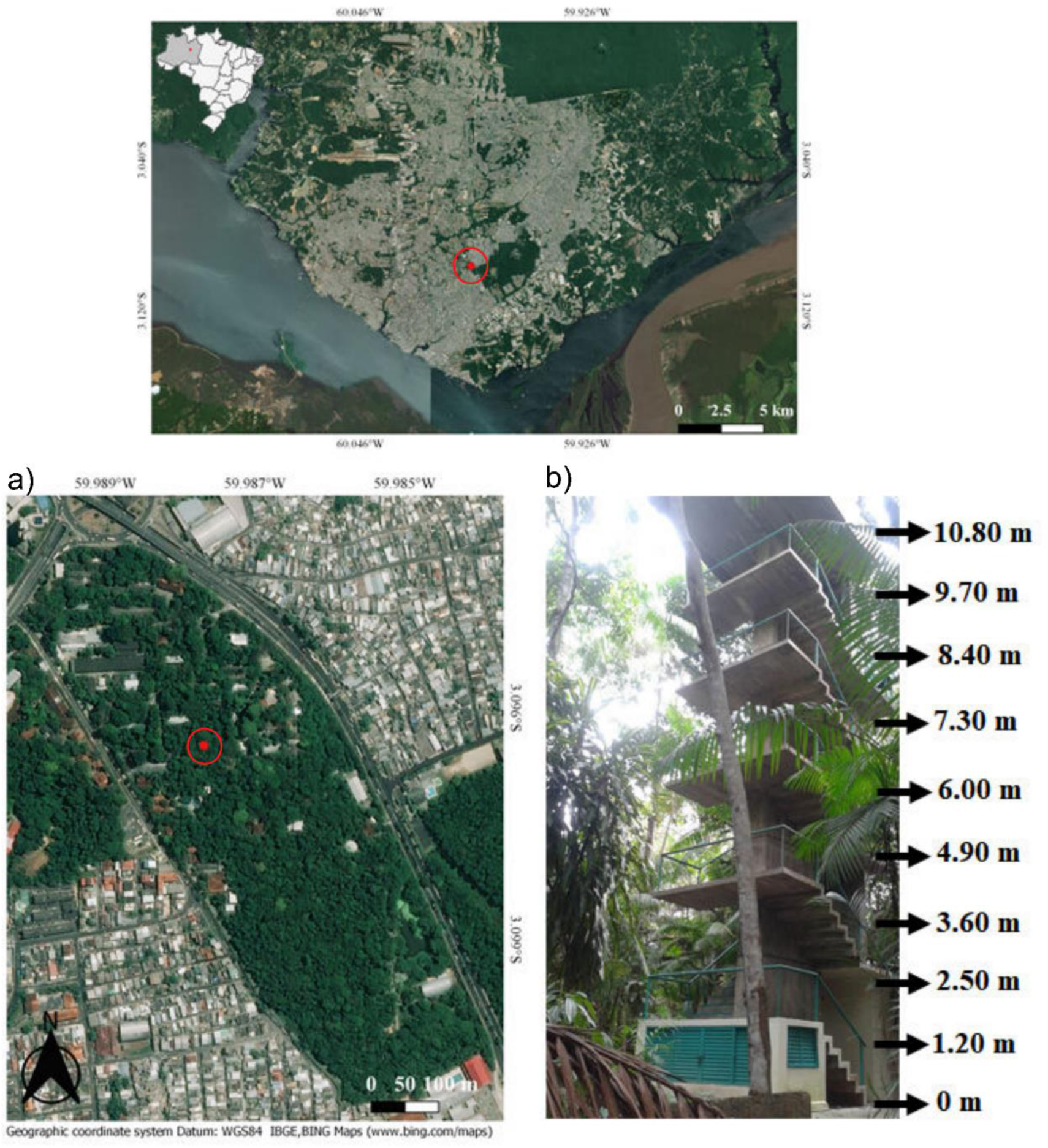
Study area located in an urban forest fragment (a), with the presence of the experimental tower and different heights (b) in Manaus, Amazonas.

### Field experiments

The experiment was carried out in a water tower with six floors, measuring 15.13 meters (m) high and 3.20 meters at the base (Figure 1b). At each height (ground – 0 m; 1.20 m; 2.50 m; 3.60 m; 4.90 m; 6 m; 7.30 m; 8.40 m; 9.70 m; and 10.8 m) three oviposition traps of the ovitrap type were installed (Fay and Eliason 1966; Depoli *et al*. 2016; Silva *et al*. 2018), with a distance of 1.50 m between each one, totaling 30 traps for each sampling period.

The traps contained 300 mL of grass infusion (1.2%) (attractant solution) and monitoring was performed daily during 10 weeks of sampling, five during the dry season (Amazon summer – September and October 2015) and another five in the rainy season (Amazon winter – March and April 2016). The traps were installed in positions protected from the sun and rain throughout the entire length of the tower, and their straws were collected and replaced with new ones every seven days. The eggs contained in the straws were sent to INPA’s Malaria and Dengue Laboratory, where quantification was carried out using a ZEISS Stemi 2000 50x stereoscopic microscope. The sampling design regarding obtaining the attractant solution and arrangement of traps at the sampling site followed that described by Silva *et al*. (2018) and Rossi-da-Silva *et al*. (2021).

### Data analysis

Temperature (°C), relative air humidity (%), and rainfall (mm) values were obtained from the automatic meteorological station installed in Manaus (A101), in the meteorological database for teaching and research (BD MAP) of the National Institute of Meteorology, located approximately 100 m from the study site (INMET, 2020).

After quantifying the collected eggs, the Ovitrap Positivity Index - OPI (OPI = Number of positive traps / Number of traps examined) x 100 and the Egg Density Index - EDI (EDI = Number of eggs / Number of positive traps) (Gomes, 1998) were calculated. The OPI was used to verify the percentage of eggs laid by *Aedes* spp. in the traps used in each collection week, considering the two seasons of the year (dry - summer; rainy - winter). The EDI was used to analyze the egg density obtained in each trap throughout the sampling period, in each season of the year.

The data were subjected to the Shapiro-Wilk normality test to verify data distribution. Subsequently, the paired t-test was performed to compare differences in egg abundance and height classes (low: 0 – 2.5m, medium: 3.6 – 6m, and high: 7.3 – 10.8m) in distinct seasonal periods (dry and rainy). Analysis of variance (two-way ANOVA) was used to compare the groups (height, season, and class height) in relation to the abundance of *Aedes* eggs. Tukey’s Post Hoc multiple comparison test was used to identify the difference in the average number of eggs x height x season, and between height classes (p=<0.05). Multiple linear regression was used to verify the relationship between the dependent/response variable (eggs) and the independent/explanatory variables (rainfall, temperature, relative humidity). All analyses were performed in R (R Development Core Team 2020).

## RESULTS

A total of 35,152 eggs were collected in the ovitraps, 18,777 in the dry season and 16,375 in the rainy season. There was no significant difference in the abundance of *Aedes* eggs between the seasonal periods analyzed (t=1.308, df=49, p=0.196). At all evaluated heights, high quantities of *Aedes* spp. eggs were obtained, however the total number showed a small decrease from a height of 3.60m in the dry season and 4.90m in the rainy season (Table 1). Considering a total of 191 pairwise comparisons at different heights and seasonal periods, twelve (12) showed significant differences in the mean number of eggs (p<0.05), as shown in Table 2.

**Table 1.** Abundance of *Aedes* spp. eggs. at different heights during five weeks of sampling, in the two seasonal periods (dry and rainy) in Manaus/Amazonas.

| Heights<br>(m) | Dry |  |  |  | Rainy |  |  |  |
| --- | --- | --- | --- | --- | --- | --- | --- | --- |
|  | Min. | Max. | Mean + SD | Total | Min. | Max. | Mean + SD | Total |
| 0 | 24 | 588 | 192 ± 142 | 2891 | 9 | 366 | 83 ± 86 | 1243 |
| 1.20 | 55 | 617 | 263 ± 143 | 3956 | 32 | 292 | 164 ± 73 | 2169 |
| 2.50 | 52 | 477 | 211 ± 121 | 3175 | 50 | 230 | 141 ± 56 | 2069 |
| 3.60 | 8 | 216 | 95 ± 66 | 1431 | 49 | 341 | 143 ± 73 | 2025 |
| 4.90 | 0 | 228 | 101 ± 65 | 1521 | 28 | 173 | 88 ± 44 | 1206 |
| 6.00 | 0 | 436 | 86 ± 113 | 1300 | 9 | 336 | 129 ± 87 | 1862 |
| 7.30 | 0 | 419 | 127 ± 116 | 1910 | 5 | 194 | 66 ± 58 | 925 |
| 8.40 | 0 | 275 | 78 ± 71 | 1174 | 35 | 208 | 96 ± 49 | 1414 |
| 9.70 | 1 | 421 | 106 ± 117 | 1488 | 58 | 152 | 89 ± 26 | 1274 |
| 10.80 | 0 | 232 | 46 ± 57 | 693 | 32 | 235 | 88 ± 53 | 1290 |

\*SD: Standard deviation.

**Table 2.**
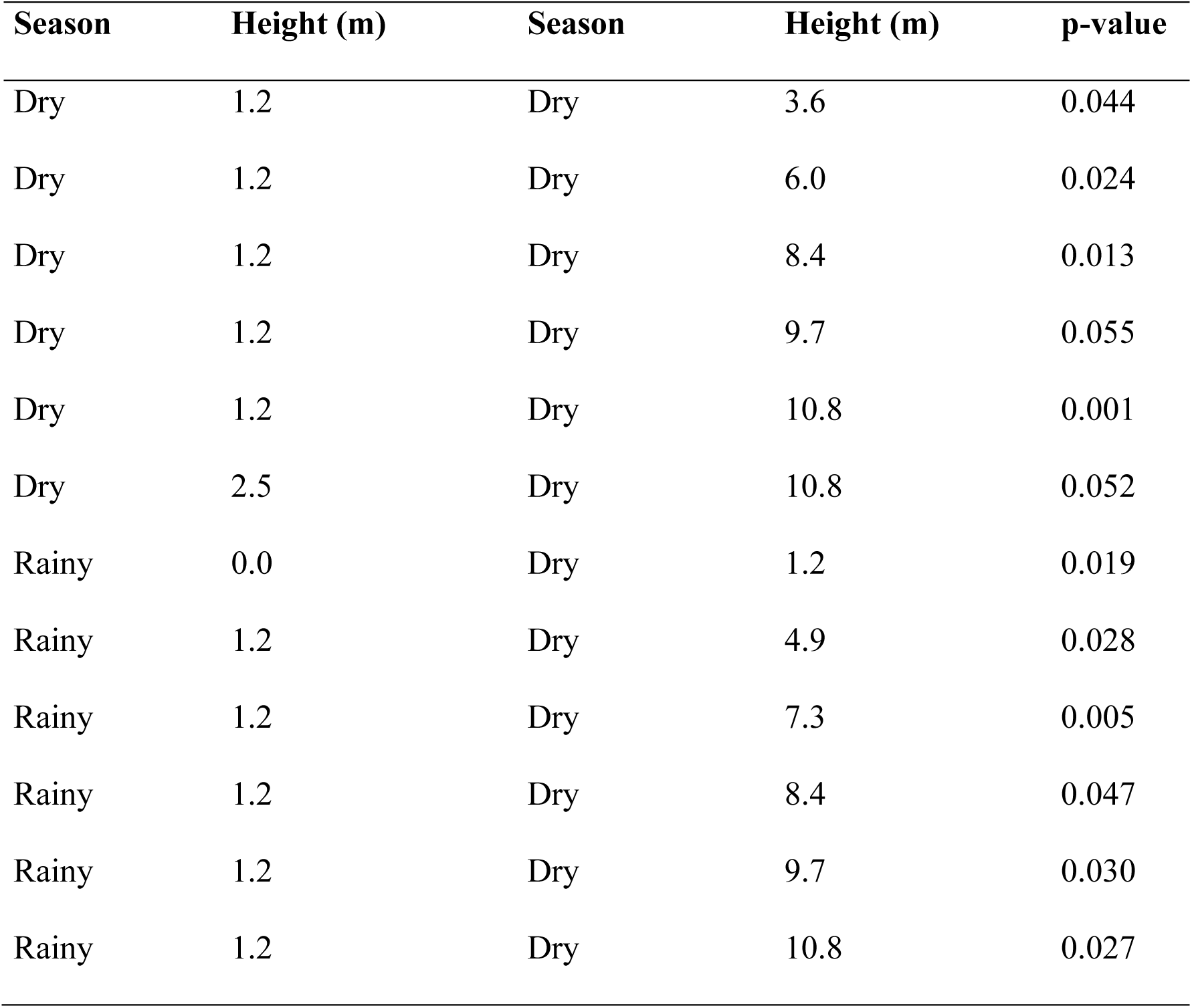
Significant results when comparing egg means between different heights and seasonal periods in Manaus, Amazonas.

A significant difference was observed in the abundance of eggs between the dry and rainy periods at low heights (0 to 1.2 meters) (df=20.9, p.adj=0.029), however, no difference was observed in other comparisons of the medium (3.6 to 6 m) and high height classes (7.3 to 10.8 m), respectively (df=23.3, p.adj=0.735; df=27.8, p.adj=1). Differences were also identified between low and high heights (df=44.7, p.adj=0.000) and low and medium heights (df=47.7, p.adj=0.007) (Figure 2).

**Figure 2.**
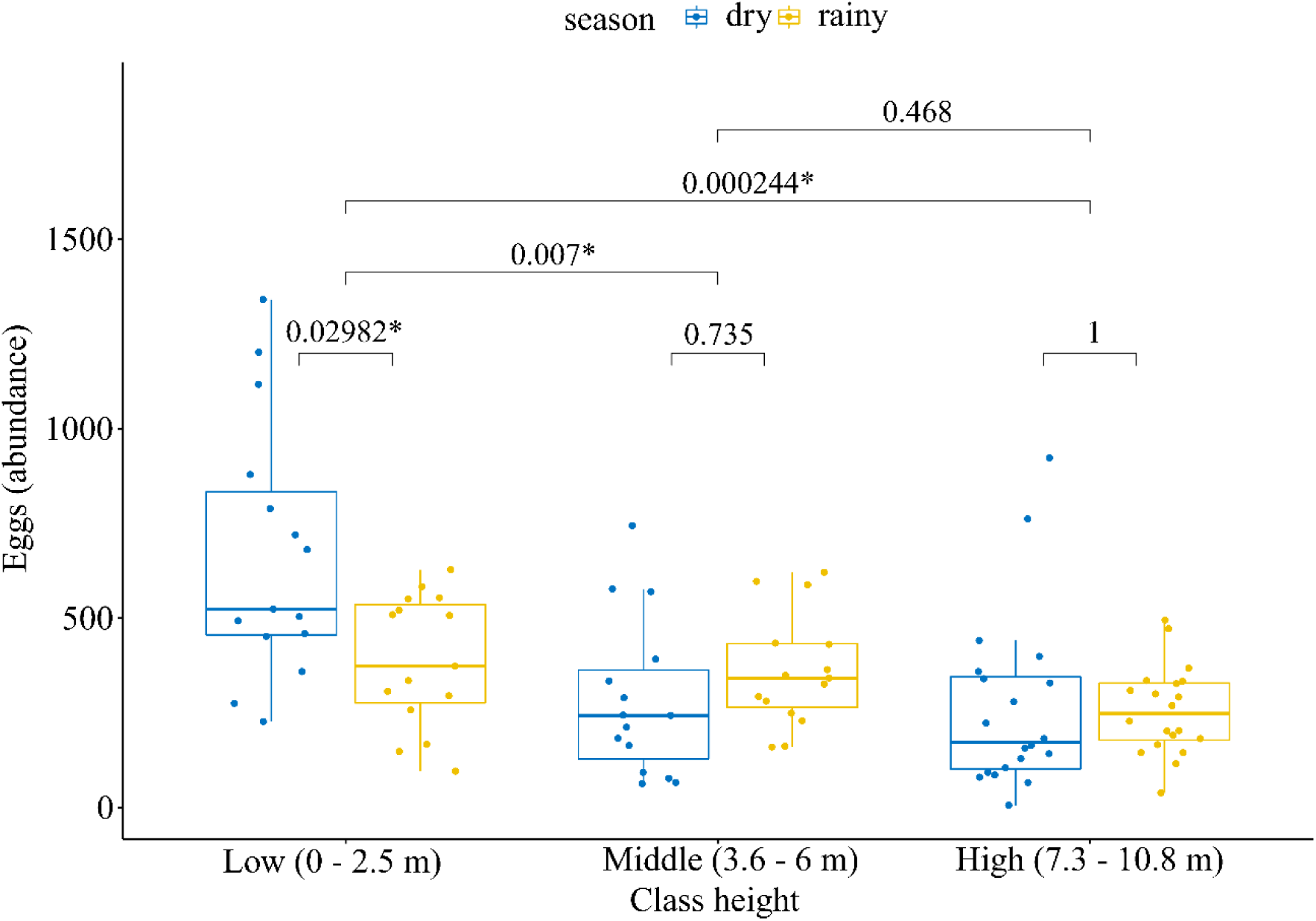
Comparisons of the average number of eggs between the dry and rainy periods in each height category and between height classes.

The OPI in the dry period was constant and with the presence of eggs throughout the sampling period. However, 100% positivity was not observed at the following heights: 4.90 m, 6.00 m, 7.30 m, 8.40 m, 10.8 m. In relation to the EDI results, the highest egg density values were observed in the first three heights of the tower (ground – 0 m; 1.20 m; 2.50 m) and these density values reduced as the height increased, with lower density at a height of 10.8 m (Table 3). On the other hand, during the rainy season, the OPI showed 100% positivity at all heights, but egg density was higher at the following heights: 1.20 m; 2.50 m; 3.60 m and the lowest value was found at a height of 7.30 m (Table 3).

**Table 3.**
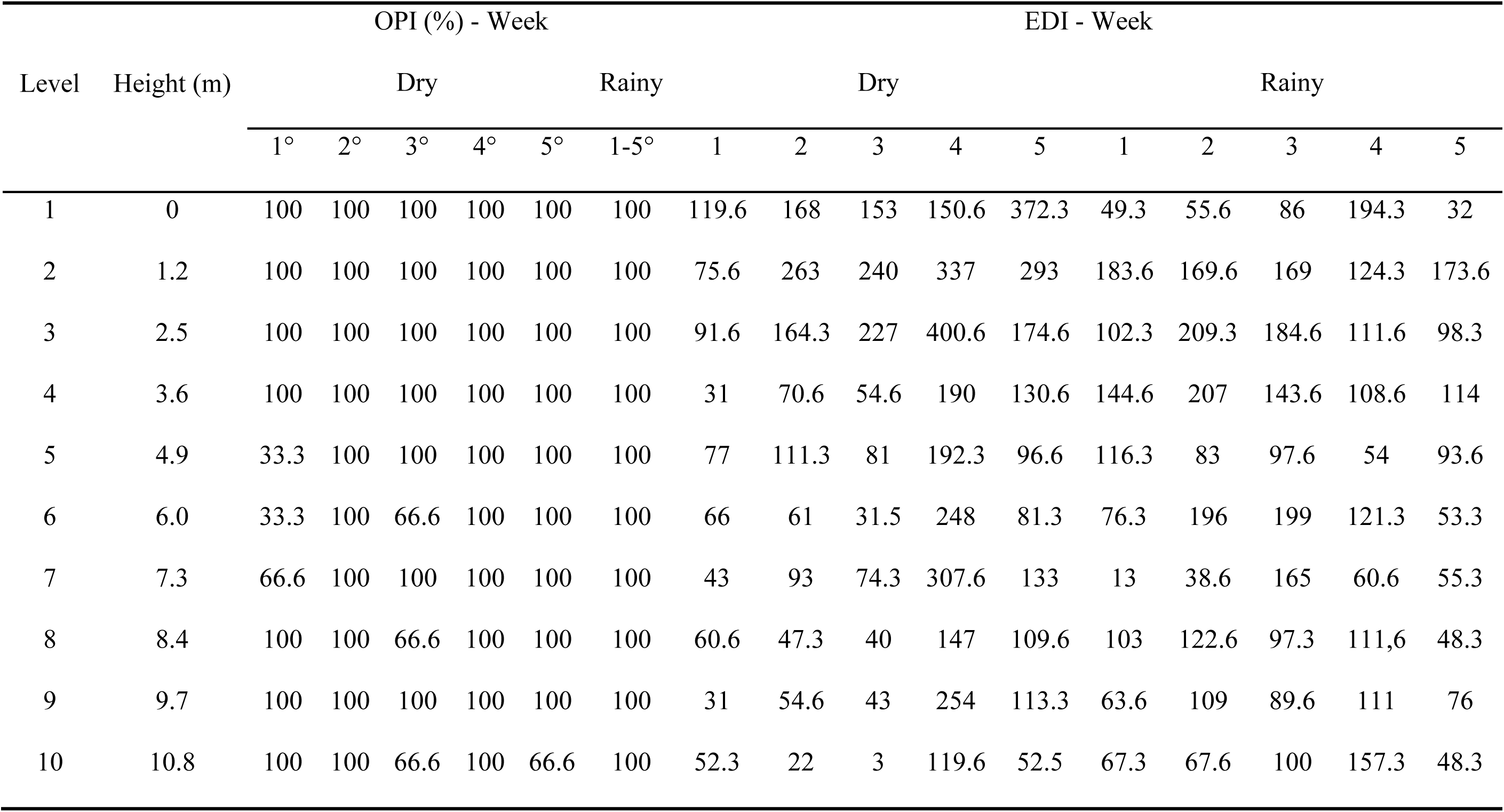
Ovitrap Positivity Index (OPI) and Egg Density Index (EDI) obtained for five consecutive weeks at different heights in the dry and rainy season in Manaus/Amazonas.

In the dry period, the temperature ranged from 29.0 to 33.7°C, with a mean of 31.7°C ± 1.1. Relative air humidity averaged 64.1% ± 9.2, ranging from 51.0 to 75.8%. The mean precipitation observed was 0.7 mm ± 2.9, with a variation from 0.0 to 15.8 mm during this period (Figure 3). In the rainy season, the average temperature was 27.7 °C ± 1.1, varying from 25.4 to 30.0°C, presenting lower temperatures compared to the dry period. Relative air humidity ranged from 70.3 to 94.5% and averaged 81.7% ± 8.6. The mean precipitation observed was 11.1 mm ± 16.0, with values ranging from 0.0 to 72.8 mm, considered the period with the highest amount of rainfall in the region (Figure 3).

**Figure 3.**
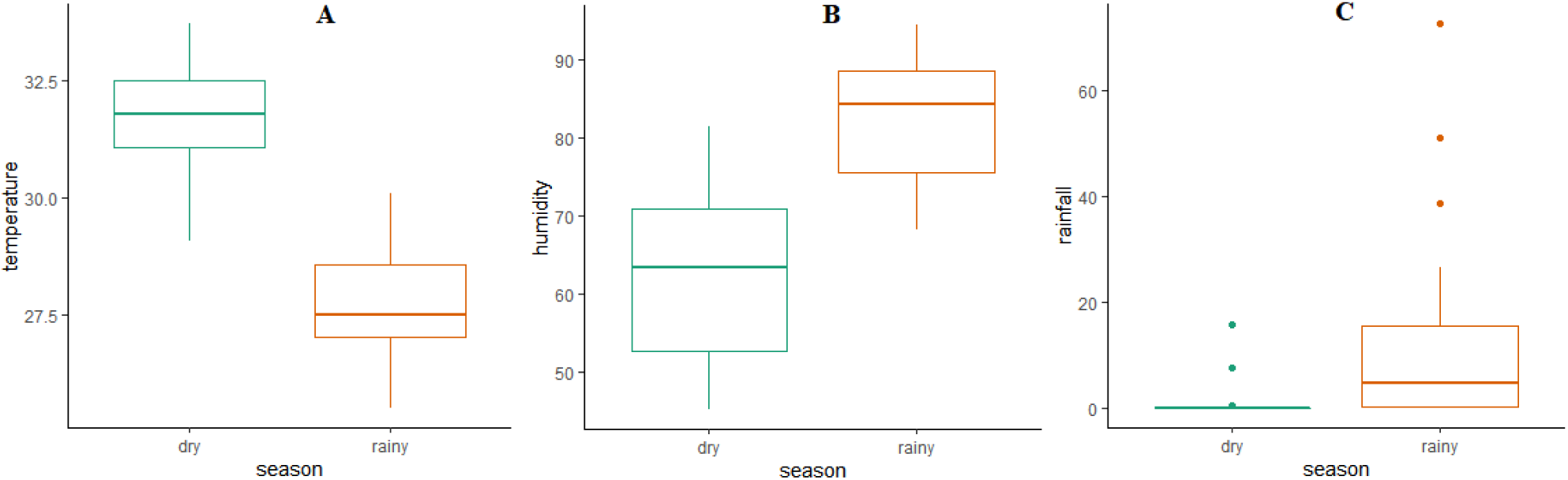
Temporal variation of temperature (A), relative humidity (B), and rainfall (C) in the two seasonal periods in Manaus, Amazonas.

The model generated in the multiple regression analysis showed that relative humidity explained 34% of the variation in *Aedes* eggs (R^2^ = 0.341, p= 0.036) in the dry period. Rainfall and temperature were not significant in this same period (R^2^ = 0.121, p= 0.199; R^2^ = 0.172, p= 0.069). During the rainy season, the variables relative air humidity (R^2^ = 0.109, p= 0.112), rainfall (R^2^ = 0.066, p= 0.377), and temperature (R^2^ = 0.081, p= 0.218), also did not influence the abundance of eggs.

## Discussion

Monitoring mosquito vectors of pathogens is essential to detect where these insects proliferate and to map cases of arboviruses (Pautasso et al. 2013; Osório et al. 2014; Medeiros et al. 2018; Connelly 2019). The data obtained in this study demonstrate that entomological surveillance can be carried out using oviposition traps (ovitraps), which are practical and effective tools for detecting the presence and abundance of *Aedes* eggs within a few days (Depoli et al. 2016; Silva et al. 2018; Nascimento et al. 2020; Silva et al. 2021).

In reference to the positivity and abundance of eggs observed in the present study, it was found that the values obtained in the seasons (dry and rainy) can be explained by the high density of mosquitoes present in the sampling area. These results corroborate the data observed by Silva et al. (2018), who installed ovitraps associated with different control agents on INPA Campuses I and II and collected 7,684 *Aedes* eggs during five weeks of the rainy season. More recently, Silva et al. (2021) using ovitraps, collected around 3,773 *Aedes* eggs on Campus I of INPA during the five-week period of the rainy season, demonstrating a high abundance of eggs in this location.

INPA Campus I has native and/or secondary forest close to the urban environment, favoring populations of *A. aegypti* and *A. albopictus*, especially the latter species, as observed by Silva et al. (2018) and Silva et al. (2021). The area contains several buildings with the constant movement of people in and out of the properties, in addition to a wide availability of breeding sites distributed throughout the fragment. These factors associated with the city’s high temperatures and recurrent rains from November to June (INMET, 2021), corresponding to the second collection period carried out in the current study, provide a favorable environment for the proliferation of these mosquitoes.

This composition of the abundance and presence of mosquitoes in urban forest fragments, such as the prevalence of *A. albopictus*, can elucidate the potential paths for the spread of arboviruses between urban and wild perimeters, which receive the constant movement of people and animals (Montagner et al. 2018; Hendy et al. 2020). It is noteworthy that the city of Manaus is constituted as a mosaic of areas with fragments of forest reserves, made up of buildings with high movement of people and animals inside and outside the properties, presenting suitable conditions for different species of mosquitoes, mainly for *A. albopictus*. This species has great medical importance, as it can infect and transmit more than 20 arboviruses, including the Chikungunya virus, representing a bridge vector in the transmission of pathogens from wild habitats to the urban environment (Ferreira-de-Lima et al. al. 2020). In the laboratory, it has vector capacity for more than 20 arboviruses (Vega-Rúa et al. 2014; Data on the total number of eggs obtained in each sampling period (dry and rainy) carried out in this study, provide evidence of the efficiency of ovitrap traps as a substrate for oviposition. These are designed to facilitate the process of oviposition and adhesion of *Aedes* spp. eggs in the field, however, when associated with an attractant solution based on grass infusion, they become more attractive for females to lay eggs (Depoli et al. 2016; Silva et al. 2018; Silva et al. 2021). This is important, as in the current study ovitraps containing grass infusion were effective for monitoring oviposition and vertical dispersal of *Aedes* spp. in a tower with different heights.

The tower chosen for this study reflects the standards of vertical housing, of the multi-story building type, built in large urban centers and close to the presence of a green area. Therefore, the study area appears to be a complete ecosystem that provides an ecological niche with fundamental biotic and abiotic components for *A. aegypti* and *A. albopictus*. As these mosquitoes deposit their eggs in multiple locations (skip oviposition), having breeding sites in different vertical strata facilitates oviposition at higher heights (Abreu et al. 2015). These same components are easily found in vertical housing, including the presence of humans, plants, and pets in apartments, while abiotic factors are related to temperature and humidity (Jayathilake et al., 2015). Collectively, this conjunction of factors provides ideal conditions for mosquito reproduction, such as a blood meal, egg-laying containers, water for larval and pupal stages, as well as a resting place for adults at various elevations in high-rise apartments (Lau et al., 2013).

The adaptive quality of egg laying by the Aedes mosquito above its flight line is demonstrated by the presence of eggs on all floors of the study tower. This characteristic is discussed in the works of Tinker (1974) and Chadee (2004), suggesting that the vertical movement of *A. aegypti* above ground level may result from insecticidal pressure on adult insects and removal of breeding sites at ground level.

In the current work, eggs of *Aedes* spp. were found more frequently at lower heights (0 to 2.5 meters), however, oviposition, even with small quantities, was also observed at higher heights, indicating that the species is capable of reproducing at any level of a residential building and that the distance in relation to the height of the ground is not an impediment to egg laying. This finding is in line with the results obtained by Roslan et al. (2013). Wan-Norafikah et al. (2010) emphasize the decrease in the population of *Aedes* mosquitoes with the height of the floors, being highest on the ground floor.

According to the data obtained in the current research, significant differences were identified in the average number of eggs at different heights, in different seasonal periods, mainly at lower heights. However, the data obtained do not corroborate the results of Obenauer et al. (2009), who detected no noticeable difference in egg collection between suburban habitats with heights of 1 and 6 m, however, a greater number of eggs were collected in ovitraps placed in wild habitats at a height of 1 m. These results are comparable to the current study, and those demonstrated by Jayathilake et al. (2015), who pointed out indices with differences in the average number of larvae per ovitrap, especially at heights of 2, 3, and 6 m in relation to the ground floor of the building.

Data from Chadee (2004) suggest that vertical dispersal and a high abundance of *A. aegypti* may occur due to biotic and abiotic components found in an ecological niche, providing blood, and breeding and resting sites in high-rise residential buildings, which present a building structure similar to that considered in this study. In addition to these, and other biotic and abiotic factors verified in the work of Lau et al. (2013) and Jayathilake et al. (2015), a general reason for this vertical dispersal of *Aedes* mosquitoes can be suggested to be the abundance of the population of these vectors in colonized habitats, whether residential or peri-urban. The percentage of forest cover and the seasonal period are indicated as important predictors for the mosquito community in general, with a strong relationship with the vector species (Arcos et al., 2021).

Considering the data from the OPI and EDI indices obtained in this study, it was found that the presence and density of *Aedes* eggs was high at different heights and in each season. This result can be corroborated with the results obtained by Silva et al. (2018), who found that the positivity (OPI) and density (EDI) of mosquito eggs was high on INPA Campuses I and II, over five weeks of sampling. The study site is a highly altered area, with the presence of buildings, people, animals, breeding sites (natural and artificial), and native and/or secondary vegetation, providing excellent conditions for the permanence and proliferation of mosquitoes of the genus *Aedes*, mainly when associated with the city’s climate, which presents abundant rain and high temperature and humidity values.

The distribution of the *Aedes* mosquito by vertical dispersal of its eggs directly causes the spread of dengue fever and other arboviruses if the local population is not immune to the circulating arboviruses. Abiotic factors, such as temperature, humidity, and rainfall have led to the presence of this vector in several urban environments around the world (WHO, 2019). With the exception of relative air humidity, which showed an influence on egg abundance in the dry period, the other environmental variables did not present a significant effect in this study. Although there is a definite effect of meteorological parameters on *Aedes* behavior, the results of Jayathilake et al. (2015), also described no significant difference in these parameters for the vertical distribution of *Aedes* mosquitoes in their study.

According to Degallier (2010), climatic factors have a direct influence on the biology of mosquitoes, since high relative humidity favors young mosquitoes, but is harmful to old mosquitoes. Average and minimum temperatures positively influence the mortality of adult and old mosquitoes respectively. Therefore, the characteristics of Manaus, such as the proximity of forest environments, with two well-defined climatic seasons (dry and rainy) and high temperatures and humidity, favor the proliferation and survival of *Aedes* spp. According to Lima et al. (2021) and Arcos et al. (2021), seasonality and forest cover directly and indirectly influence the mosquito larval community, mainly in the peri-urban area of Manaus.

Furthermore, another striking characteristic of the eggs of this vector is the ability to remain viable for long periods when water is scarce, thus resisting until the next period of rain. This period of quiescence allows for the indirect and involuntary transport of eggs to other regions by human action (Mayilsamy, 2019). Climatic factors must be taken into account to develop more sustainable and efficient prevention strategies against dengue (Junxiong & Yee-Sin, 2015).

## Final considerations

With the expansion of the urban perimeter and vertical growth in urban areas of large cities such as Manaus, we believe that updated information on the vertical dispersal of mosquitoes of the genus *Aedes* in this habitat is necessary in order to verify the concentration, mobility, and presence of breeding sites of this and other potential vectors for public health, and should be included in entomological monitoring and control. It is noteworthy that lower heights provide a greater abundance of *Aedes* eggs, however, this is not a limiting factor for colonization at other heights. Climatic factors, such as relative air humidity end up positively influencing the average number of eggs of these mosquito vectors in the urban area of Manaus, especially during the dry period, since the levels of this variable are very unstable during the season.

## Acknowledgements

The authors thank the team of the Laboratory of Malaria and Dengue of the National Institute of Research of the Amazon- INPA. This study was financed in part by the Brazilian National Council for Scientific and Technological Development (CNPq), Amazonian Research Foundation (FAPEAM) and Coordination for the Improvement of Higher Education Personnel – Brasil (CAPES - Financing Code 001).

## References

Ab Hamid, N., Mohd Noor, S. N., Isa, N. R., Md Rodzay, R., Bachtiar Effendi, A. M., Hafisool, A. A., … & Lee, H. L. (2020). Vertical infestation profile of *Aedes* in selected urban high-rise residences in Malaysia. Tropical medicine and infectious disease, 5(3), 114.

Abreu, F. V. S. D., Morais, M. M., Ribeiro, S. P., & Eiras, Á. E. (2015). Influence of breeding site availability on the ovOPIsition behaviour of *Aedes aegypti*. Memórias do Instituto Oswaldo Cruz, 110(5), 669–676.

Ayllón, T., Câmara, D. C. P., Morone, F. C., Gonçalves, L. D. S., Saito Monteiro de Barros, F., Brasil, P., … & Honório, N. A. (2018). Dispersion and oviposition of *Aedes albopictus* in a Brazilian slum: Initial evidence of Asian tiger mosquito domiciliation in urban environments. PLoS One, 13(4), e0195014.

Araújo, R. A. F., Uchôa, N. M., & Alves, J. M. B. (2019). Influência de Variáveis Meteorológicas na Prevalência das Doenças Transmitidas pelo Mosquito *Aedes aegypti*. Revista Brasileira de Meteorologia, 34(3), 439–447.

Arcos, A.N.; Valente-Neto, F.; Ferreira, F.A.S.; Bolzan, F. P.; Cunha, H. B.; Tadei, W. P.; Hughes, R. M.; Roque, F. O. (2021). Seasonality modulates the direct and indirect influences of forest cover on larval anopheline assemblages in western Amazônia. Scientific Reports, 11, 12721. 10.1038/s41598-021-92217-9

Bentley MD, Day JF 1989. Chemical ecology and behavioral aspects of mosquito ovOPIsition. Annual Review of Entomology 34: 401–421.

Bergero, Paula E.; Ruggerio, Carlos A.; Lombardo, Ruben; Schweigmann, Nicolás J.; Solari, Hernán G. Dispersal of *Aedes aegypti*: Field study in temperate areas using a novel method. Journal of Vector Borne Diseases 50(3):p 163–170, Jul–Sep 2013.

Bharati, M., & Saha, D. (2021). Status de resistência a inseticidas e mecanismos bioquímicos envolvEDIs em mosquitos *Aedes*: uma revisão de escopo. Asian Pacific Journal of Tropical Medicine, 14 (2), 52.

Bonizzoni, M.; Gasperi, G.; Chen, X.; James, A.A. 2013. The invasive mosquito species *Aedes albopictus*: current knowledge and future perspectives. Trends in parasitology 29: 460–468.

Campbell, L.P.; Luther, C.; Moo-Llanes, D.; Ramsey, J.M.; Danis-Lozano, R.; Peterson, A.T. 2015. Climate change influences on global distributions of dengue and chikungunya virus vectors. Philosophical Transactions of the Royal Society B: Biological Sciences 370: 20140135.

Carvalho, F.D.; Moreira, L.A. 2017. Why is *Aedes aegypt*i Linnaeus so successful as a species?. Neotropical Entomology 46: 243–55.

Chadee, D. D. (2004). Observations on the seasonal prevalence and vertical distribution patterns of ovOPIsition by *Aedes aegypti* (L.) (Diptera: Culicidae) in urban high-rise apartments in Trinidad, West Indies. Journal of vector ecology: journal of the Society for Vector Ecology, 29(2), 323–330.

Clements, A.N. 1992. The Biology of Mosquitoes: development, nutrition and reproduction. Chapman & Hall, London, 1: 509.

Connelly, R. (2019). Highlights of Medical Entomology 2018: The Importance of Sustainable Surveillance of Vectors and Vector-Borne Pathogens. Journal of medical entomology, 56(5), 1183–1187.

Consoli, R. A. G. B., de Oliveira, R. L. Principais mosquitos de importância sanitária no Brasil. SciELO-Editora FIOCRUZ, 1994.

Couret J, Dotson E, Benedict MQ. Temperature, larval diet, and density effects on development rate and survival of *Aedes aegypti* (Diptera: Culicidae). PLoS One. 2014; 9(2): 1–9.

Custódio, J. M. D. O., Nogueira, L. M. S., Souza, D. A., Fernandes, M. F., Oshiro, E. T., Oliveira, E. F. D., … & Oliveira, A. G. D. (2019). Abiotic factors and population dynamic of *Aedes aegypti* and *Aedes albopictus* in an endemic area of dengue in Brazil. Revista do Instituto de Medicina Tropical de São Paulo, 61.

Da Silva, V. C., Scherer, P. O., Falcão, S. S., Alencar, J., Cunha, S. P., Rodrigues, I. M., & Pinheiro, N. L. (2006). Diversidade de criadouros e tOPIs de imóveis frequentados por *Aedes albopictus* e *Aedes aegypti*. Revista de Saúde Pública, 40(6), 1106–1111.

Degallier, N. Impactos climáticos sobre a transmissão da dengue no nordeste do Brasil. Projeto Clima do Atlântico Tropical e Impactos sobre o Nordeste, FUNCEME: Fortaleza, p. 331-337, 2010.

Deng, S.Q.; Yang, X.; Wei, Y.; Chen, J.T.; Wang, X.J.; Peng, H.J. A review on dengue vaccine development. Vaccines 2020, 8, 63.

Depoli, P. A. C., Zequi, J. A. C., Nascimento, K. L. C., & Lopes, J. (2016). Eficácia de ovitrampas com diferentes atrativos na vigilância e controle de *Aedes*. EntomoBrasilis, 9(1), 51–55.

Dickens, B.L.; Sun, H.; Jit, M.; Cook, A.R.; Carrasco, L.R. (2018). Determining environmental and anthropogenic factors which explain the global distribution of *Aedes aegypti* and *Ae. albopictus*. BMJ global health 3: e000801.

Fay, R.W. & Eliason, D.A. A preferred ovOPIsition site as surveillance method for *Aedes aegypti*. Mosquito News, v.26, n.4, p.531-535, 1966.

Ferreira-de-Lima, V. H., Câmara, D. C. P., Honorio, N. A., & Lima-Camara, T. N. (2020). The Asian tiger mosquito in Brazil: Observations on biology and ecological interactions since its first detection in 1986. Acta Tropica, 205, 105386.

Forattini, O.P. (2002). CulicEDIlogia Médica. São Paulo: Editora USP, São Paulo-SP. 860p.

Goddard, J., Moraru, G. M., Mcinnis, S. J., Portugal, J. S., Yee, D. A., Deerman, J. H., & Varnado, W. C. (2017). A statewide survey for container-breeding mosquitoes in Mississippi. Journal of the American Mosquito Control Association, 33(3), 229–232.

Gomes, A. de C. Medidas dos níveis de infestação urbana para *Aedes (Stegomyia) aegypti* e *Aedes (Stegomyia) albopictus* em programas de Vigilância Entomológica. Informes Epidemiológico do SUS, v.7, n.3, p.49-57, 1998.

Gonzalez PV, González Audino PA, Masuh HM. OvOPIsition Behavior in *Aedes aegypti* and Aedes albopictus (Diptera: Culicidae) in Response to the Presence of Heterospecific and Conspecific Larvae. J Med Entomol. 2016 Mar;53(2):268–72. doi: 10.1093/jme/tjv189. PMID: 26634825.

González, M. A., Rodríguez-Sosa, M. A., Vásquez-Bautista, Y. E., del Carmen Rosario, E., Durán-Tiburcio, J. C., & Alarcón-Elbal, P. M. (2020). A survey of tire-breeding mosquitoes (Diptera: Culicidae) in the Dominican Republic: Considerations about a pressing issue. Biomédica, 40(3), 507.

Harbach, R.E. 2023. Mosquito taxonomic inventory. (http://mosquito-taxonomicinventory.info/). Acesso em 22 de dezembro de 2023.

Harrington LC, Ponlawat A, Edman JD, Scott TW, Vermeylen F 2008. Influence of container size, location and time of day on ovOPIsition patterns of the dengue vector, Aedes aegypti, in Thai- land. Vector-Borne Zoonotic Disease 8: 415–424.

Hasanuddin, I., Miyagi, I., Toma, T. & Kamimura, K. (1997). Breeding habitats of *Aedes aegypti* (L.) and *Aedes albopictus* (Skuse) in villages of Barru, South Sulawesi, Indonesia. Southeast Asian Journal of Tropical Medicine and Public Health 28 (4): 844–850.

Hashim, N. A., Ahmad, A. H., Talib, A., Athaillah, F., & Krishnan, K. T. (2018). Co- breeding association of *Aedes albopictus* (Skuse) and *Aedes aegypti* (Linnaeus) (Diptera: Culicidae) in relation to location and container size. Tropical life sciences research, 29(1), 213.

Hendy A, Hernandez-Acosta E, Valério D, Mendonça C, Costa ER, Júnior JTA, Assunção FP, Scarpassa VM, Gordo M, Fé NF, Buenemann M, de Lacerda MVG, Hanley KA, Vasilakis N. The vertical stratification of potential bridge vectors of mosquito-borne viruses in a central Amazonian forest bordering Manaus, Brazil. Sci Rep. 2020 Oct 26;10(1):18254. doi: 10.1038/s41598-020-75178-3. PMID: 33106507; PMCID: PMC7589505.

Honório, N. A., Câmara, D. C. P., Calvet, G. A., Brasil, P. Chikungunya: uma arbovirose em estabelecimento e expansão no Brasil. Cadernos de Saúde Pública, 2015, v. 31, p. 906-908.

INMET - National Institute of Meteorology. Meteorological Database for Teaching and Research, 2020. Available in: http://www.inmet.gov.br/portal/index.php?r=bdmep/bdmep.

Instituto Brasileiro de Geografia e Estatística - IBGE. Disponível em: https://cidades.ibge.gov.br/brasil/am/manaus/panorama. Acesso em: 22 de fevereiro de 2021.

Iwamura, T., Guzman-Holst, A. & Murray, K. A. Accelerating invasion potential of disease vector *Aedes aegypti* under climate change. Nature Communication 11, 2130 (2020). 10.1038/s41467-020-16010-4

Jayathilake T. A.; Wickramasinghe M. B.; de Silva B. G. OvOPIsition and vertical dispersal of *Aedes* mosquitoes in multiple storey buildings in Colombo district, Sri Lanka. Jounal of Vector Borne Disease 2015; 52(3): 245–51.

Junxiong P, Yee-Sin L. Clustering, climate and dengue transmission. Expert Rev Anti Infect Ther. 2015;13(6):731–740.

Kamal, M.; Kenawy, M.A.; Rady, M.H.; Khaled, A.S.; Samy, A.M. (2018). Mapping the global potential distributions of two arboviral vectors *Aedes aegypti* and *Ae. albopictus* under changing climate. PloS one 13: e0210122.

Kato, F.; Ishida, Y.; Oishi, S.; Fujii, N.; Watanabe, S.; Vasudevan, S.G.; Tajima, S.; Takasaki, T.; Suzuki, Y.; Ichiyama, K.;, et al (2016). Novel antiviral activity of bromocriptine against dengue virus replication. Antivir. Res., 131, 141–147.

Kraemer, M.U.; Sinka, M.E.; Duda, K.A.; Mylne, A.Q.; Shearer, F.M.; Barker, C.M.; Moore, C.G.; Carvalho, R.G.; Coelho, G.E.; Van Bortel, W.;, et al. (2015). The global distribution of the arbovirus vectors Aedes aegypti and Ae. albopictus. elife 4: e08347.

Lau, K. W., Chen, C. D., Lee, H. L., Izzul, A. A., Asri-Isa, M., Zulfadli, M., & Sofian-Azirun, M. (2013). Vertical distribution of *Aedes* mosquitoes in multiple storey buildings in Selangor and Kuala Lumpur, Malaysia. Tropical biomedicine, 30(1), 36–45.

Lemos, L. S. M.; Costa, R. C. Bacias Hidrográficas em Manaus (2005 – 2015). In: Costa, R. C. (Org.) Riscos, fragilidades & problemas ambientais urbanos em Manaus. Manaus: Editora INPA, 2017, pp. 129–135.

Leta, S.; Beyene, T.J.; De Clercq, E.M.; Amenu, K.; Kraemer, M.U.; Revie, C.W. (2018). Global risk mapping for major diseases transmitted by *Aedes aegypti* and *Aedes albopictus*. International Journal of Infectious Diseases 67: 25–35.

Liew, C. C. F. C., & Curtis, C. F. (2004). Horizontal and vertical dispersal of dengue vector mosquitoes, *Aedes aegypti* and *Aedes albopictus*, in Singapore. Medical and Veterinary Entomology, 18(4), 351–360.

Lima, G. R., Santos, E. V., Arcos, A. N., Lima, C. A. P., Tadei, W. P., & Castro Simões, R. (2021). Abundância larval de *Anopheles* em criadouros artificiais na zona leste de Manaus, Amazonas. South American Journal of Basic Education, Technical and Technological, 8(1), 35–47.

Lourenço-de-Oliveira, R. (2015). Biologia e comportamento do vetor. Valle, D.; Pimenta, D.N.; Cunha, R.V.; organizadores. Dengue teorias e práticas. Rio de Janeiro: Fiocruz, 76–92.

Marcondes, C.B.; Ximenes, M.D.F.F.D. (2016). Zika virus in Brazil and the danger of infestation by *Aedes* (*Stegomyia*) mosquitoes. Revista da Sociedade Brasileira de Medicina Tropical 49: 4–10.

Mayilsamy M. Extremely Long Viability of *Aedes aegypti* (Diptera: Culicidae) Eggs Stored Under Normal Room Condition, *Journal of Medical Entomology*, Volume 56, Issue 3, May 2019, Pages 878–880.

Medeiros, A. S., Costa, D. M., Branco, M. S., Sousa, D. M., Monteiro, J. D., Galvão, S. P., … & Araújo, J. M. (2018). Dengue virus in *Aedes aegypti* and *Aedes albopictus* in urban areas in the state of Rio Grande do Norte, Brazil: Importance of virological and entomological surveillance. PLoS One, 13(3), e0194108.

Montagner, F. R. G.; Silva, O. S.; Jahnke, S. M. Mosquito species occurrence in association with landscape composition in green urban áreas. Braz. J. Biol. 2018, vol. 78, no. 2, pp. 233–239.

Nascimento KLC, da Silva JFM, Zequi JAC, Lopes J. Comparison Between Larval Survey Index and Positive Ovitrap Index in the Evaluation of Populations of *Aedes* (*Stegomyia*) *aegypti* (Linnaeus, 1762) North of Paraná, Brazil. Environmental Health Insights. 2020 Jan 6;14:1178630219886570. doi: 10.1177/1178630219886570. PMID: 31933523; PMCID: PMC6945453.

Navarro-Silva MA, Marques FA, Duque LE, Jonny E (2009). Review of semiochemicals that mediate the ovOPIsition of mosquitoes: a possible sustainable tool for the control and monitoring of Culicidae. Revista Brasileira de Entomologia 53: 1–6.

Obenauer P. J.; Kaufman P. E.; Allan A. S.; Kline D. L. (2009). Ovitrampas com iscas por infusão para pesquisar as preferências de altura ovOPIsicional de mosquitos que habitam contêineres em dois habitats da Flórida, Journal of Medical Entomology, Volume 46, Issue 6, Pages 1507–1513.

Osório, H. C., Zé-Zé, L., Amaro, F., & Alves, M. J. (2014). Mosquito surveillance for prevention and control of emerging mosquito-borne diseases in Portugal—2008– 2014. International journal of environmental research and public health, 11(11), 11583–11596.

Pautasso, A., Desiato, R., Bertolini, S., Vitale, N., Radaelli, M. C., Mancini, M., … & Casalone, C. (2013). Mosquito Surveillance in Northwestern I taly to Monitor the Occurrence of Tropical Vector-Borne Diseases. Transboundary and emerging diseases, 60, 154–161.

Pontes GO; Silva WR; Ferreira FAZ; Roque RA; Tadei WP; Zequi JAC; Fé NF; Freitas AB; Vieira SS; Justiniano SCB. Dimorfismo sexual de pupas de *Aedes (Stegomyia) aegypti* Linnaeus, 1762 e *Aedes (Stegomyia) albopictus* (Skuse, 1894) (Diptera: Culicidae). Scientia Amazônia. V9, n2. B1–B8, 2020.

Powell, J.R.; Gloria-Soria, A.; Kotsakiozi, P. (2018). Recent history of *Aedes aegypti*: Vector genomics and epidemiology records. BioScience 68: 854–860.

R Development Core Team. (2020). R: A language and environment for statistical computing. R foundation for statistical computing, Vienna, Austria. (Available from: http://www.R-project.org/).

Rossi da Silva K, Ribeiro da Silva W, Silva BP, Arcos AN, da Silva Ferreira FA, Soares-da-Silva J, et al. (2021) New traps for the capture of *Aedes aegypti* (Linnaeus) and *Aedes albopictus* (Skuse) (Diptera: Culicidae) eggs and adults. PLoS Negl Trop Dis 15(4): e0008813. 10.1371/journal.pntd.0008813.

Roslan MA, Shafie A, Ngui R, Lim YAL, Sulaiman WYW. Vertical infestation of the dengue vectors *Aedes aegypti* and *Aedes albopictus* in apartments in Kuala Lumpur, Malaysia. Journal of the American Mosquito Control Association 2013; 29(4): 328–36.

Sanoussi, A. F., Loko, L. Y., Ahissou, H., Adjahi, A. K., Orobiyi, A., Agré, A. P., … & Sanni, A. (2015). Diversity, physicochemical and technological characterization of elite Cassava (Manihot esculenta Crantz) Cultivars of Bantè, a District of Central Benin. The Scientific World Journal, 2015.

Santos, I. C. D. S., Braga, C., de Souza, W. V., de Oliveira, A. L. S., & Regis, L. N. (2020). The influence of meteorological variables on the ovOPIsition dynamics of *Aedes aegypti* (Diptera: Culicidae) in four environmentally distinct areas in northeast Brazil. Memórias do Instituto Oswaldo Cruz, 115.

Silva, W. R. D., Soares-da-Silva, J., Ferreira, F. A. D. S., Rodrigues, I. B., Tadei, W. P., & Zequi, J. A. C. (2018). OvOPIsition of *Aedes aegypti* Linnaeus, 1762 and *Aedes albopictus* Skuse, 1894 (Diptera: Culicidae) under laboratory and field conditions using ovitraps associated to different control agents, Manaus, Amazonas, Brazil. Revista Brasileira de Entomologia, 62(4), 304–310.

Soares-da-Silva, J., Ibiapina, S. S., Bezerra, J. M. T., Tadei, W. P., & Pinheiro, V. C. S. (2012). Variation in *Aedes aegypti* (Linnaeus) (Diptera, Culicidae) infestation in artificial containers in Caxias, state of Maranhão, Brazil. Revista da Sociedade Brasileira de Medicina Tropical, 45(2), 174–179.

Soares-Pinheiro, V.C.; Dasso-Pinheiro, W.; Trindade-Bezerra, J.M.; Tadei, W.P. (2017). Eggs viability of *Aedes aegypti* Linnaeus (Diptera, Culicidae) under different environmental and storage conditions in Manaus, Amazonas, Brazil. Brazilian Journal of Biology 77: 396–401.

Tadei, W. P.; Pinto, R. C.; Santos, J. M. M.; Rodrigues, I. B.; Rafael, M. S.; Albuquerque, B. C.; Fialho, R. R. Cal e Cloro no Controle do Mosquito da Dengue. O Biólogo, v. 1, p. 14-15, 2011.

Tham, A. S. (2000). Surveillance of mosquitoes. Mosquitoes and mosquito-borne diseases: biology, surveillance, control, personal and public protection measures. Kuala Lumpur, Malaysia: Academy of Sciences Malaysia. *p*, 167–183.

Tinker, M. E. (1974). *Aedes aegypti* larval habitats in Surinam. Bulletin of the Pan American Health Organization (PAHO); 8 (4).

Vega-Rúa, A.; Zouache, K.; Girod, R.; Failloux, A.B.; Lourenço-de-Oliveira, R. (2014). High level of vector competence of *Aedes aegypti* and *Aedes albopictus* from ten American countries as a crucial factor in the spread of Chikungunya virus. Journal of Virology 88: 6294–6306.

Wan-Norafikah O, Nazni WA, Noramisa S, Shafa’ar-Ko’ohar S, Azirol-Hisham A, Nor- Hafizah, R, et al, Vertical dispersal of *Aedes* (*Stegomyia*) spp. in high-rise apartments in Putrajaya, Malaysia. Trop Biomed 2010; 27(3): 662–7.

WHO. Dengue and Severe Dengue. Available online: https://www.who.int/news-room/fact-sheets/detail/dengue-and-severe-dengue (accessed on 10 January 2021).

WHO. WHO Region of the AMERICAS Records Highest Number of Dengue Cases in History; Cases Spike in Other Regions. Available online: https://www.who.int/news-room/detail/21-11-2019-who-region-of-the-americas-records-highest-number-of-dengue-cases-in-history-cases-spike-in-other-regions (accessed on 15 January 2021).

WRBU. 2023. Walter Reed Biosystematics Unit. Systematic catalog of Culicidae. Washington, USA. Disponível em: http://www.mosquitocatalog.org/default.aspx?pgID=2. Acesso em 12/06/2023.

